# Optical Fiber-Assisted Bioprinting Enables Freeform Printing of Cell-Laden and Turbid Hydrogel Resins

**DOI:** 10.64898/2026.05.28.728387

**Authors:** Maximilian Pfeiffle, Alessandro Cianciosi, Sebastian Beusink, Tomasz Jüngst

**Affiliations:** Department of Functional Materials in Medicine and Dentistry, Institute of Biofabrication and Functional Materials, University of Würzburg, Pleicherwall 2, 97070 Würzburg, Germany; AO Research Institute Davos, Davos Platz, Switzerland; Department of Orthopedics, Regenerative Medicine Center Utrecht, University Medical Center Utrecht, Heidelberglaan 100, 3584 CX Utrecht, The Netherlands

**Keywords:** Biofabrication, Bioprinting, Light-Based, GelMA, PEGDA, OFAB (Optical Fiber-Assisted Bioprinting)

## Abstract

Optical fiber-assisted printing (OFAP) was recently introduced as a straightforward light-based platform for the spatially controlled photopolymerization of hydrogel-based resins. Here, we extend this concept toward optical fiber-assisted bioprinting (OFAB) by processing cell-laden GelMA- and GelMA/PEGDA-based bioresins in a freeform embedded printing configuration. The system relies on a 405 nm LED-coupled optical fiber mounted on an automated 3D motion platform, enabling localized photocrosslinking directly within a resin bath. First, GelMA and GelMA/PEGDA formulations containing LAP and tartrazine were screened to evaluate the influence of light intensity, printing velocity, and material composition at the line width and curing depth. Single-line features with widths down to 70 ± 20 µm were obtained under optimized conditions, while more robust printing conditions yielded reproducible features in the range of 200–300 µm. Photorheological and rotational rheology measurements confirmed that the formulations provide both thermoresponsive support during printing and photocrosslinked stability after processing. The incorporation of L929 cells demonstrated high cytocompatibility for GelMA and GelMA/PEGDA 6000 Da formulations, with viabilities above 90% after 7 days for selected printed constructs. Importantly, increasing the cell concentration up to 1 × 10^7^ cells mL^−1^ did not prevent printing and reduced the extent of overcuring, suggesting that cell-induced turbidity can improve spatial confinement of polymerization in OFAB. Finally, a customized OFAB printer was developed to enable temperature-controlled processing and the fabrication of centimeter-scale 3D structures, including cell-laden constructs. Overall, this work establishes OFAB as an accessible and modular bioprinting strategy for cell-laden and optically turbid hydrogel resins, complementing existing light-based biofabrication approaches.

## 1. Introduction

Biofabrication is a new and quickly evolving research field^[1]^ with the ambitioned goal to revolutionize modern medicine by printing human tissue and organs,^[2]^ thus solving issues like the shortage of organ donations^[3],[4]^ which is greatly increased given the higher life expectancy^[2]^ and reducing or replacing in vivo testing.^[3],[4]^ Although the field is far away from meeting these goals, it has made great progress and the velocity of progress depends on the availability and affordability of equipment amongst other factors.^[1]^ Thus, it is important to not only research clinical applications from a biological aspect but also elaborating new and improved cytocompatible fabrication technologies. An important category of suitable automated fabrication technologies for biofabrication is Bioprinting. It uses concepts of Additive manufacturing (AM), where 3D structures are created by the addition of material in a layer-by-layer fashion. In contrast to traditional AM technologies, bioprinting is aiming for hierarchical 3D structures mimicking living tissue by directly incorporating biologically active substances like cells.^[5],[6],[7]^ Currently there is a variety of different technologies applied in biofabrication including extrusion and light-based technologies.

Extrusion-based bioprinting relies on a cell containing bioink being extruded through a nozzle, which requires a precise control over the material rheology. A high viscosity in combination with shear thinning properties helps the printed constructs to stay in shape after the print,^[8]^ however it also exposes cells to mechanical forces during extrusion which might harm them or alter their properties.^[9]^ Moreover, due to the layer-by-layer fashion most extrusion-based printers dispense inks, convoluted structures are difficult to produce^[10]^ and the lack of self-supporting properties of the bioinks can lead to the collapse of whole structures.^[11]^ Thus, there is a tradeoff between good rheological and mechanical properties for printability while maintaining ideal cell environment. This can limit the mimicking of biological features.^[8]^ To overcome some of these issues and print in a more freeform manner, hydrogel-based bioinks can be extruded into a support bath, which serves as a temporary support and may be dissolved later.^[9],[12]^ Using a method called freeform reversible embedding of suspended hydrogels (FRESH), these support baths also enable the print of low viscosity bioinks by using a shear-thinning fluid bed which is solid at low shear stress and becomes more viscos at high shear stress, enabling a self-healing of the support bath.^[2],[13]^ The maximal resolution however is limited with extrusion-based methods to about 100 μm due to nozzle size and maximal stress cells can tolerate.^[9],[10]^

Light-based methods on the other hand, can achieve hight print resolutions of down to 100 nm^[14]^ while giving an unprecedented degree of freedom in fabricating convoluted structures^[15]^ often using photocrosslinkable hydrogel precursors acting as resins.^[16]^ These include biologically derived materials like Gelatin methacryloyl (GelMA), which is a crosslinkable version of gelatin and the gold-standard for light-based biofabrication due to its ease of production, cost-effectiveness^[17]^ and its inherent signalling molecules for cell adhesion.^[18]^ Is has excellent biocompatibility and thermoresponsive properties, causing it to gel at low temperatures.^[19]^ Often also synthetic materials like Polyethylene glycol diacrylate (PEGDA) are used, since they are more stable and provide better shape fidelity. PEGDA is known for its non-cytotoxicity (at higher molecular weights) and non-immunogenicity.^[18]^ Biologically derived and synthetic hydrogel materials can be referred to as bioresins when including cells and biomaterial resins when suited for cell seeding after the print following the definition by Groll et. al..^[16]^ Some of the most important light-based methods in biofabrication are stereolithography (SLA)/digital light processing (DLP), two-photon polymerization (2PP) and Volumetric bioprinting (VBP).^[20]^ SLA and its related version DLP are vat-based polymerisation techniques producing structures in layer-by-layer manner by laser scanning in case of SLA or projecting single layers in case of DLP in a bottom up or top-down manner. They reach resolutions of down to 10 μm.^[21]^ However, they are limited in the inks they can use since very soft inks, which can be preferential for biological applications, often deform or collapse during printing. ^[10],[21]^ For this reason, a combination of PEGDA and GelMA was used by Shu-Yung Chang at al. to combine the stability of PEGDA with the biocompatibility of GelMA in a DLP printing setup.^[22]^ A two-component ink composed of GelMA and long-chain PEGDA was also DLP printed by He et. al. to print a corneal implant supporting cell adhesion, proliferation and migration.^[23]^ DLP resolution is limited by the projector’s pixel size and optical blur. In contrast, 2PP uses nonlinear absorption, so polymerization occurs only at the laser focus where the intensity is highest,^[24]^ enabling resolutions down to ∼100 nm.^[25],[24],[26]^This is achieved using femtosecond lasers, where polymerization happens only when two photons are simultaneously absorbed by the photoinitiator at the focal point.^[25]^ While 2PPs resolution is unrivalled, the low throughput presents a challenge.^[27]^ Additionally, it is expensive, limiting its availability for a bigger range of users.^[20]^ Since transparent bioresins are used,^[28]^ the addition of cells might pose additional problems due to light scattering. Another light-based method that is much faster and can fabricate centimetre sized structures within a few seconds is volumetric bioprinting.^[29]^ It builds up an object layer-less by projecting subsequent tomographic patterns inside a rotating vial with resolutions of 40 μm to 100 μm.^[30],[31]^ Materials with higher viscosity are often used to reduce sedimentation and transparency of the resin are critical parameter affecting the printing resolution of the method.^[17]^ However, many resin formulations that include cells might lead to light scattering,^[30],[31]^ decreasing resolution. In addition, VBP is also limited by the vial size used as with bigger vials, absorption increases and print quality drops.

Optical fiber-assisted printing (OFAP) was recently introduced by our team as a light-based fabrication platform in which a LED-coupled optical fiber is integrated into an automated motion system to locally photocrosslink photosensitive resins. In contrast to projection-based vat photopolymerization, the light source is moved directly across or within the resin, allowing spatially confined curing without requiring projection optics or a fixed build area. The first demonstration of OFAP focused on biomaterial resin patterning and proof-of-concept freeform 3D printing, highlighting the possibility of adapting light intensity, fiber velocity, and fiber position during fabrication.

However, the translation of OFAP toward true bioprinting requires several additional challenges to be addressed. First, the resin formulation must simultaneously provide cytocompatibility, sufficient support during embedded printing, and post-printing stability. Second, the presence of cells introduces light scattering and turbidity, which are often detrimental to light-based bioprinting processes but may affect optical-fiber-based curing differently because polymerization is initiated locally at the fiber tip. Third, the printing workflow must be adapted to process cell-laden hydrogels under temperature-controlled and reproducible conditions.

Here, we address these challenges by developing optical fiber-assisted bioprinting (OFAB) as a cell-compatible extension of OFAP. GelMA and GelMA/PEGDA formulations were selected because GelMA provides thermoresponsive support and cell-adhesive motifs, while PEGDA can improve post-crosslinking stability. The formulations were characterized in terms of printing resolution, rheological behavior, cytocompatibility, and printability at increasing cell densities. Finally, a customized OFAB printer was implemented to demonstrate the fabrication of complex 3D hydrogel structures, including cell-laden constructs.

## 2. Results and Discussion

### 2.1 Resolution analysis of single lines with OFAB

To establish the processing window of OFAB, we first investigated how the main dose-controlling parameters, namely light intensity, and fiber translation velocity, affect the geometry of single printed lines. Single-line printing was selected as a reductionist assay because it provides direct information on the smallest achievable features and on the anisotropy between lateral line width and axial curing depth. Three hydrogel formulations were compared to decouple the effects of material composition, optical turbidity, and post-crosslinking stability on OFAB resolution.

The formulations were selected to represent biologically relevant soft hydrogel systems rather than fully optimized engineering resins. GelMA was used because of its widespread application in biofabrication, thermoresponsive gelation, and cell-adhesive motifs, whereas PEGDA was incorporated to improve post-crosslinking stability and construct handling. This material selection allowed us to evaluate the balance between cytocompatibility, support-bath behavior, and printing fidelity.

When printing soft hydrogels with OFAB, it is helpful to have a strong support structure to keep soft lines with small diameters intact during the print, especially since they constantly detach from the optical fiber tip, which puts a strain on them. Therefore, GelMA based bioinks are a good material choice since the thermo-gelling properties of GelMA can be used as a support structure. To do so, the formulations are cooled to 5°C before the print. Afterwards the resin can be 3D patterned. The uncrosslinked resin is removed by heating the printed structure up to 37° degrees (**Fehler! Verweisquelle konnte nicht gefunden werden**.). The resolution of an OFAB print is determined by the material composition and the system settings. While the material formulation can be chosen freely, ranging from very stiff formulations of 70% PEGDA 700 Da^[32]^ down to soft hydrogels composed of 6% GelMA, the most key factors are the concentration of photo-absorber and initiator. In this work, Lithium phenyl-2,4,6-trimethylbenzoylphosphinate (LAP) and tartrazine was used. However, while the possibility of resin formulations is broad, the system settings are limited. In order to make a reasonable assumption about the OFAB systems printing resolution range, the system specific parameters including the intensity setting and printing velocity were screened using three different soft hydrogel formulations with 6% GelMA(6G), 3% PEGDA 6000 Da and 3% GelMA (6G_3P6000) and 3% PEGDA 700 Da and 3% GelMA (3G_3P700) all including 0.15% LAP as photoinitiator and 0.015% tartrazine as photoabsorber in 254 mM HEPES as a pH buffer.

Single lines were printed with different light intensities as depicted in **Fehler! Verweisquelle konnte nicht gefunden werden**. A. Those single lines give an indication of printing resolutions when printing simple structures like cubes (**Fehler! Verweisquelle konnte nicht gefunden werden**. B, C). Since the emitted light of the optical fiber is, in the given setup, vertical to the base-plate, the line depth and width have to be viewed separately. When reducing the light Intensity from 6.8 mW/cm^2^ (LED setting of 100%) to 3.7 mW/cm^2^ (LED setting of 50%) at a velocity of 0.25 mm/s all three formulations show a continues decrease of polymerisation depth with significant differences between the highest and lowest possible setting (**Fehler! Verweisquelle konnte nicht gefunden werden**. D). The smallest intensity of 2.0 mW/cm^2^ was only able to produce stable lines with the 3G_3P700 formulation, reaching a polymerisation depth of 570 ± 70 µm. This is due to the material’s higher stability of the low molecular weight PEGDA, making the printing of smaller lines possible.^[33]^ The lowest setting able to print all formulations was 3.7 mW/cm^2^ (50%) reaching polymerisation depths down to 860 ± 120 µm for G, 680 ± 80 µm for 3G_3P6000 and 810 ± 90 µm for 3G_3P700 compared to their 1270 ± 150 µm, 1020 ± 100 µm and 1110 ± 90 µm counterparts at the highest setting. The 6G formulation preformed significantly worse than the 3G_3P6000 formulation throughout all Intensities and worse than the 3G_3P700 at the highest setting. However, the difference between the 3G_3P700 and 3G_3P6000 formulations was insignificant, which sets apart OFAB from other light-based bioprinting platforms, since the 3G_3P6000 formulation was turbid while the 3G_3P700 formulation is fully transparent (**Fehler! Verweisquelle konnte nicht gefunden werden**.). When considering the line width of the same printed structures (**Fehler! Verweisquelle konnte nicht gefunden werden**. E), the lowest expected linewidth would be 100 µm since this is the fiber diameter of the used optical fiber. 3G_3P700 reached a minimal width of 140 ± 20 µm at the 2.0 mW/cm^2^ setting. At 3.7 mW/cm^2^ 6G, 3G_3P6000 and 3G_3P700 reached the diameters of 270 ± 70 µm, 250 ± 50 µm and 230 ± 30 µm compared to 410 ± 110 µm, 370 ± 110 µm and 330 ± 30 µm at the highest setting with no significant differences within the groups. This demonstrates that the material composition has a low influence on the printed line widths.

Another system parameter affecting the light dosage is the printing velocity. To evalute its influence on resolution, the printing velocity was varied ranging from 0.1 mm/s to 0.5 mm/s at a constant intensity of 6.8 mW/cm^2^ and the impact on the line depth was analysed as shown in **Fehler! Verweisquelle konnte nicht gefunden werden**. F. The increased velocity overall reduced the line diameter significantly over all groups. The 3G_3P700 formulation reached its lowest diameter at a velocity of 1 mm/s with 250 ± 70 µm. However, fiber generation had high failure rates of 50% at these setting and is therefore not included in the figure. The highest velocity where all materials were printable at was 0.5 mm/s and resulted in depths of 860 ± 210 µm for the G, 810 ± 140 µm for the 3G_3P6000 and 870 ± 70 µm for the 3G_3P700 formulation. In-between the groups there were only a few setting that enable to generate fibers with significant differences in depth resolution, which were not systematic. The line width is also significantly decreased with the printing velocity throughout all formulations while the difference between the materials at the same velocity are not always significant (**Fehler! Verweisquelle konnte nicht gefunden werden**. G). At 0.5 mm/s, widths of 340 ± 100 µm (G), 240 ± 40 µm (P6k) and 250 ± 30 µm (P700) were printed. The lowest linewidth was produced with 1 mm/s and the 3G_3P700 formulation 70 ± 20 µm with low reproducibility, which overcame the assumed minimum of 100 µm of the fiber diameter. With those results, the OFAB system can, in terms of resolution, be placed close to extrusion printings maximum of 100 µm^[10]^ and at the lower end of Volumetric printing,^[31]^ while the printing velocity is similar to slow extrusion-based printing. Though the realistic top resolutions of OFAB might be higher than the demonstrated ones since the formulations where not optimised for resolution and the printing of single lines over a gap might yield worse results than printing a tightly knitted 3D mesh. Overall, the trends show that the printing velocity and light intensity both significantly influence the overall resolution and highlight the systems current capabilities. Furthermore, the height of the lines was always a multitude bigger than their width, making the OFAB printed single lines uneven shaped. This uneven shaping between the longitudinal or orthogonal direction of the projected light, is not exclusive to OFAB, but affects other light-based bioprinting methods like SLA/DLP when the curing times and light intensities are not optimised.^[34]^ Those well-established methods, however, have different dose tests and resolution assessments as described by Levato et. al..^[34]^ These printed lines were the first attempt of creating such a test for OFAB, however more testing and the reduction of the line depth by adding more photo absorber might be needed in the future. While there are often no significant differences in resolution at the same settings between the material groups, there were clear differences in stability.

Overall, these results show that OFAB resolution is primarily governed by the delivered optical dose, which can be tuned through light intensity and fiber velocity. While the tested formulations showed only moderate differences in lateral line width under comparable conditions, they differed in stability and reproducibility. This indicates that material selection in OFAB should not be based solely on minimum feature size, but also on the ability of the printed hydrogel to withstand detachment from the fiber tip, thermal removal of the uncrosslinked bath, and post-printing handling.

### 2.2 Photorheological analysis of the formulation

Because GelMA and GelMA/PEGDA blends serve simultaneously as resin and temporary support bath in OFAB, rheological characterization was performed to clarify their stability before, during, and after photocrosslinking. Photorheological time sweeps were used to evaluate light-induced stiffening, whereas rotational measurements were used to assess viscosity, shear-thinning behavior, and time-dependent stability under constant shear. These measurements are important because OFAB requires the uncrosslinked bath to support the printed features during fabrication while still allowing fiber movement and post-print removal.

First, a time sweep was conducted as depicted in Figure 2 A showing the storage and loss modulus of the different formulations before, during and after polymerisation.

**Figure 1:**
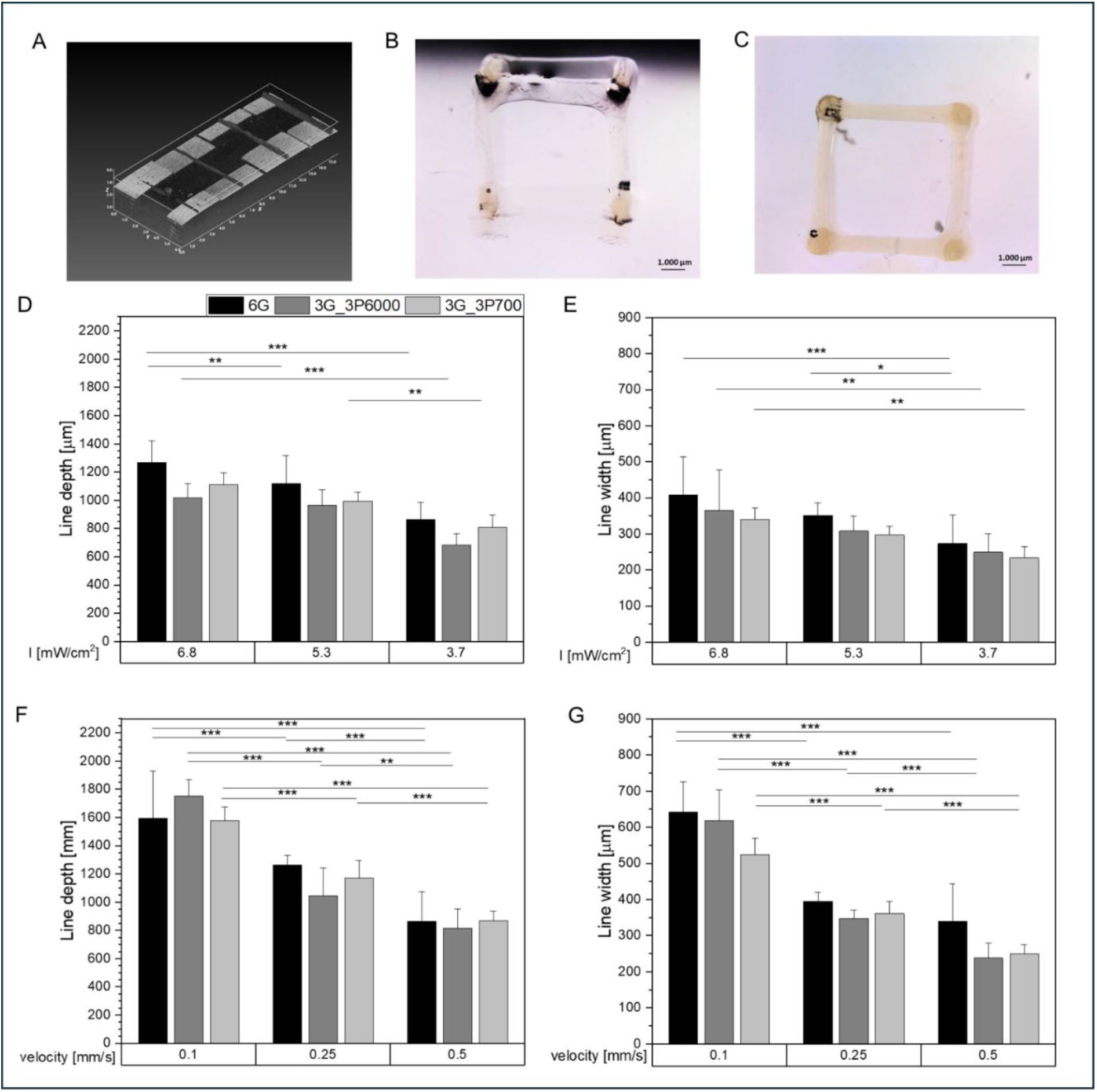
Printing resolution and characterisation of different resin compositions. (A) Depicts the OCT scan of single lines printed over a gap between two plates with different intensity settings, resulting in different widths and hights. All measurements were conducted with 3 printed lines measured over the whole gap at 5 distinct positions, resulting in an n = 15. (B) Microscope image of the side of a cube consisting of single lines, printed with 3% PEGDA 700 Da, 3% GelMA, 0.15% LAP and 0.015% tartrazine stored in water. (C) Top view of the cube from B. (D) The OCT measured polymerisation depths of single lines printed with 0.25 mm/s, dependent on the light intensity and the resin formulation. (E) The OCT measured widths (n=15) of single lines printed with 0.25 mm/s, dependent on the light intensity and the resin formulation. (F) The OCT measured polymerisation depths of single lines printed with I 6.8 mW/cm^2^, dependent on the printing velocity and the resin formulation. (G) The OCT measured line widths of single lines printed with I 6.8 mW/cm^2^, dependent on the printing velocity and the resin formulations.

**Figure 2:**
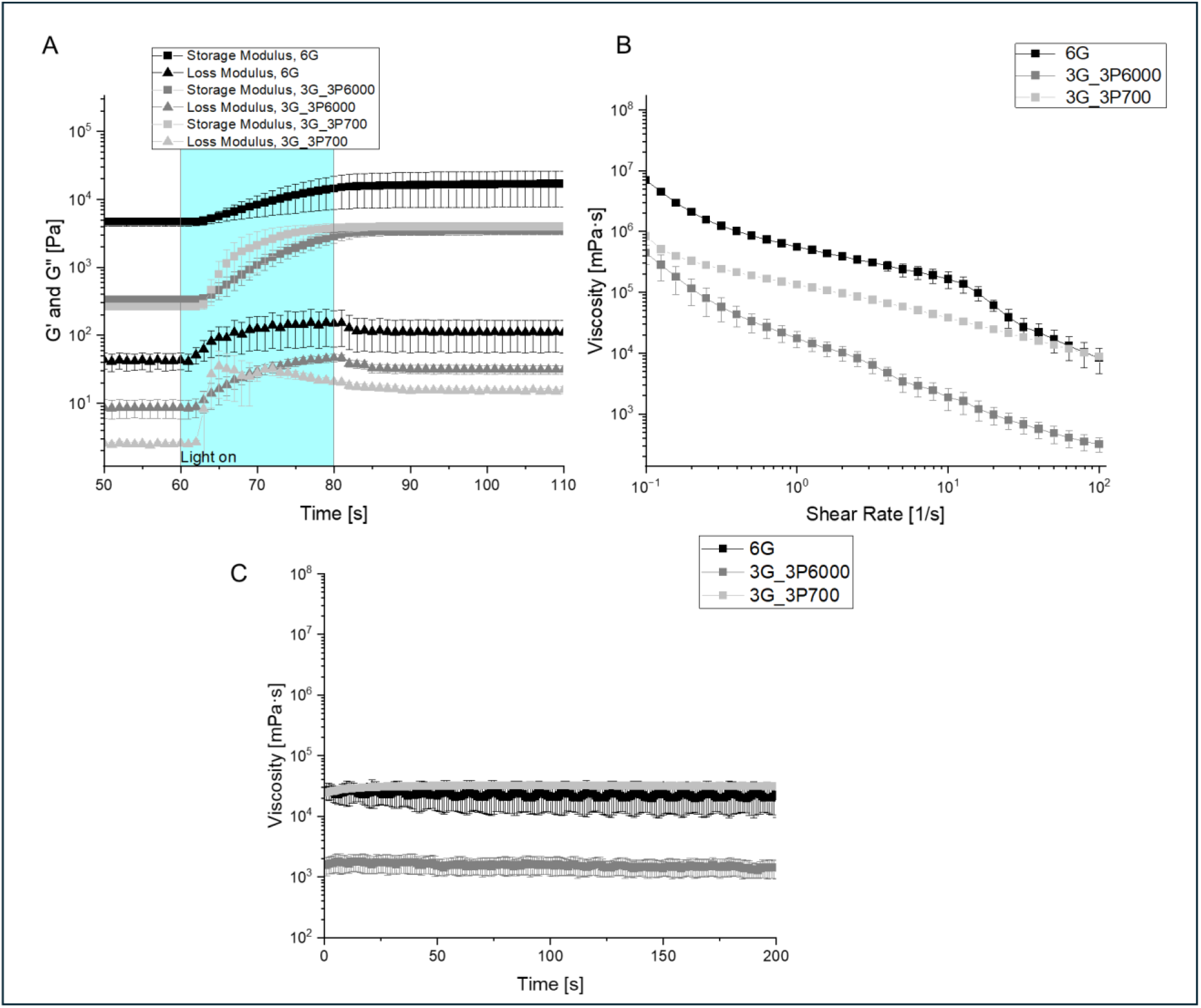
Photorheological measurements of the formulations before and during the polymerisation at 5 °C. (A) Time sweep for the different material formulations during the of a photo rheology measurement showing the change in storage and loss modulus during polymerisation between second 60 and 80. All formulations also include 0.15% LAP and 0.015% tartrazine. (B) Viscosity measured over shear rate for the different material formulations before polymerisation. (C) Time sweep at constat shear rate of 10 [1/s] for all material formulations uncross linked.

Before the polymerisation, the 6G formulation proved to be the most stable with a G’ of 4.5 x 10^3^ Pa and G’’ of 4.3 x 10^1^ Pa compared to the G’ of 3.3 x 10^2^ Pa and G’’ of 8.6 Pa of the 3G_3P6000 formulation and the G’ of 2.6 x 10^2^ Pa and G’’ of 2.5 Pa for the 3G_3P700 formulation. This is in line with the expectations, since the measurements were conducted at 5°C to mimic the printing process and the higher GelMA concentration was expected to deliver the highest stability, due to its thermo responsitivity. After 20 seconds of polymerisation, the 6G formulation maintained the highest stability with G’ of 1.6 x 10^4^ Pa and G’’ of 1.1 x 10^2^ Pa compared to P700s G’ of 3.5 x 10^3^ Pa and G’’ 1.5 x 10^1^ Pa and P6ks G’ of 4.0 x 10^3^ Pa and G’’ of 3.1 x 10^1^ Pa. The difference between G’ and G’’ of the 3G_3P700 formulation is higher compared to the 3G_3P6000 formulation, showing its increased stability and less gel like character. While the 6G formulation shows the highest stability during the print, the 3G_3P700 formulations is the most stable after the remaining support bath was washed away due to its synthetic and short monomer.^[33]^ This demonstrating that the stiffness of the support bath is not as important as the formulations stability after the crosslinking, since the 3G_3P700 formulation turned out to be more printable than the very 3G_3P6000 formulation with a similar stuff support bath and the 6G formulation. The viscosity sweep was conducted before the polymerisation (Figure 2 B), showing all formulations to be shear thinning. This supports the use of GelMA based resins as not only as resin but also self-healing support.^[11]^ While the 3G_3P700 and 6G formulations behave similar, the 3G_3P6000 formulation is more viscos than the other formulations at most shear rates before the polymerisation. In Figure 2 C the viscosity over the time period at the same shear rate of 10 1/s was measured. All materials show a constant viscosity over the measuring time, depicting that the viscosity is not dependent on the time of measurement but only of the shear rate.

In this chapter it was shown that the material with the highest GelMA concentration, provided the strongest support bath, which was within expectations, since the modulus of the support bath can be tuned with the GelMA concentration.^[35]^ In turn this suggests, that the lower printability of the material with only GelMA is not caused by the support bath stability, but by the fact that incorporation of PEGDA to the hydrogels enhances the mechanical properties,^[36]^ which however only shows its affect after melting since the crosslinked pure GelMA has still higher moduli at 5° C, compared to the blends. As described by Yue et. al. who tested different GelMA/PEGDA blends had shear thinning and self-recovery properties, all materials tested here have proven to be sheer thinning,^[33]^ which is a requirement for use as support bath,^[35]^ with the blends being more shear thinning than the pure GelMA.

Taken together, the rheological data support the dual role of the GelMA-containing formulations as both printable resin and temporary support bath. The thermoresponsive GelMA network provides initial stability at low temperature, whereas the photocrosslinked network determines the stability of the final construct after removal of the uncrosslinked material. This distinction is important for OFAB because the printed filaments must remain stable during fiber movement, detachment from the fiber tip, and post-print washing. The GelMA/PEGDA 6000 Da formulation therefore represents the most relevant compromise for bioprinting, combining cytocompatibility with improved stability compared with pure GelMA.

### 2.3 Implementation of Cells

After defining the processing window of the cell-free formulations, we next evaluated whether these materials and the OFAB process itself are compatible with cell encapsulation. L929 fibroblast-like cells were selected as a robust model cell line to decouple the biological feasibility of the process from tissue-specific differentiation effects. Cast samples were used as material controls, whereas printed samples were used to assess the additional impact of OFAB processing, including exposure to 405 nm light, handling outside standard culture conditions, and post-print washing.

In a first assay, the different polymers GelMA 6%, PEGDA 6000 Da 6% and PEGDA 700 Da 6% each with 0.15% LAP and 1×10^6^/ml L929 mouse cells were casted in silicon forms and polymerised with a 60 mW/cm^2^ 405 nm lamp for 2 minutes. A live/dead assay was conducted (Figure 3 A) to evaluate the cytocompatibility of the polymers.

**Figure 3:**
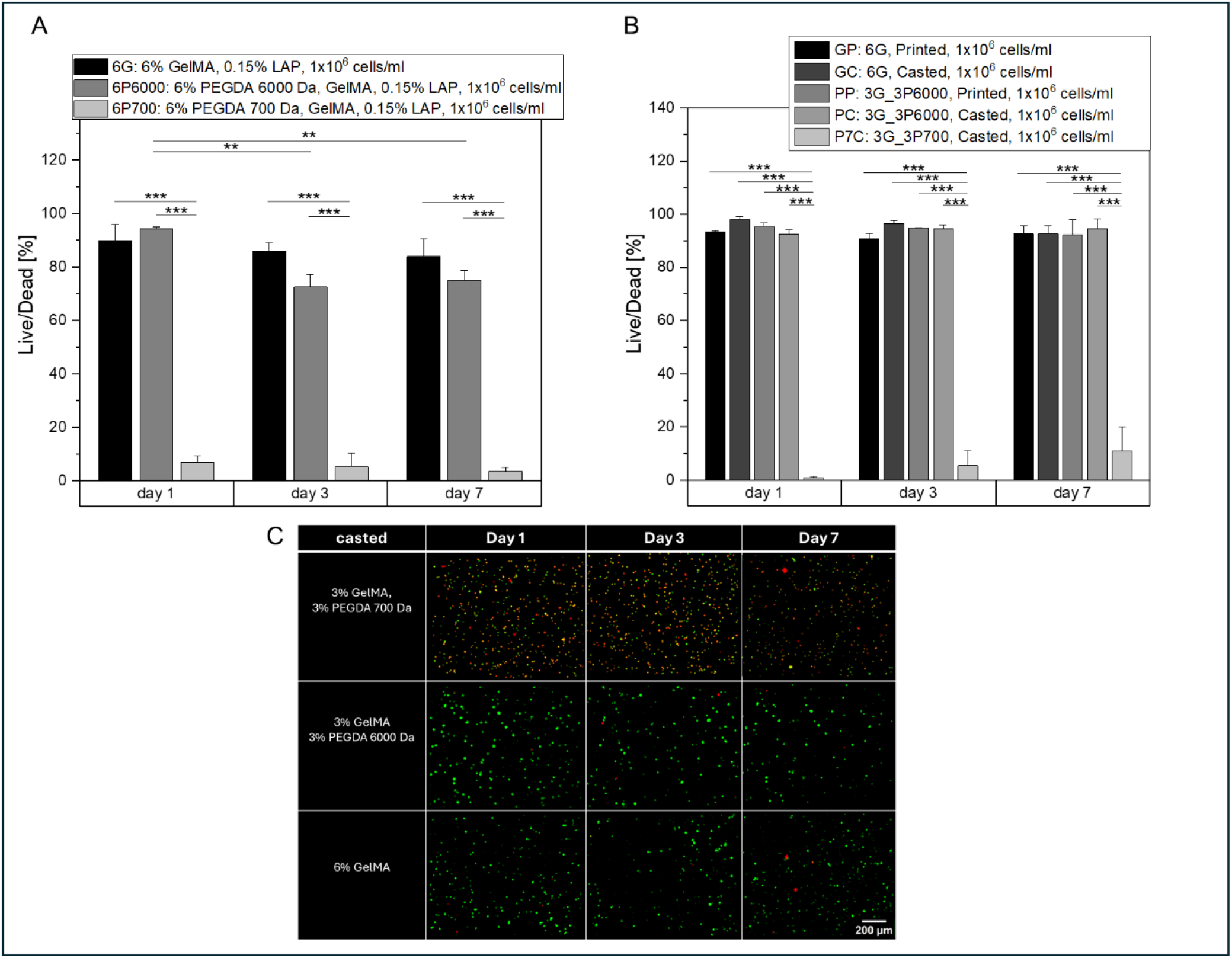
Cytocompatibility of different processes and resin formulations. (A) Live/Dead assay of 6% GelMA, 6% PEGDA 6000 Da and 6% PEGDA 700 with 0.15% LAP with 1×10^6^/ml L929 mouse cells. (B) Live/Dead assay of different resin formulations printed and casted with 1x106/ml L929 mouse cells. (C) Depiction of the live/dead assays of the different resin formulations day 1-7.

The PEGDA 700 Da based resin showed 90% of all cells had damaged membranes and proved to be unsuited for biofabrication. Although this material might be suitable as a biomaterial ink^[22]^ it cannot be used as bioink at this formulation. There was no significant difference between the pure GelMA and PEGDA 6000 Da within 7 Days, both showing high cell survival rates. However, the number of living cells significantly decreased for PEGDA 6000 Da after day 1, which might be due to the lack of cell attachment sites in the synthetic polymer.^[33]^ In a second live/dead assay the ink formulations G, 3G_3P6000 and 3G_3P700 were casted with 1×10^6^/ml L929 cells and 6G and 3G_3P6000 were additionally printed with OFAB. The experiment showed that the 3G_3P700 was not cytocompatible, since all cells were damaged. However, the 6G and 3G_3P6000 showed excellent cytocompatibility with cell survival rates over 90% at all days independent of whether they were printed or casted. This proves that OFAB is as cytocompatible as casting at the tested conditions, despite the fact that the cells have spent 30-60 min outside ideal conditions. An exemplary comparison between the resins in shown in Figure 3C. It is clear, that all cells that were in contact with PEGDA 700 Da were damaged, while the other cells retained an intact membrane.

These results indicate that OFAB is cytocompatible under the tested conditions when suitable bioresin formulations are used. Importantly, the cytocompatibility was formulation-dependent: GelMA and GelMA/PEGDA 6000 Da supported high viability, whereas PEGDA 700 Da-containing formulations were not suitable for direct cell encapsulation despite their favorable printability. This distinction identifies 3G_3P6000, rather than 3G_3P700, as the most relevant formulation for subsequent bioprinting studies.

### 2.4 Influence of cells on the OFAB print

Because OFAB initiates polymerization locally at the optical fiber tip, we hypothesized that optically turbid and highly cellularized formulations could be processed differently from conventional projection-based light bioprinting methods. We therefore investigated how increasing cell density affects both cell viability and line geometry using the GelMA/PEGDA 6000 Da formulation.

Figure 4 A depicts that with increasing cell density, the bioresins get less transparent and scatter more light. This typically effects the resolution of light-based bioprinting methods.^[34],[37]^ The light scattering is stronger, the more cells are introduced, therefore the influence of the concentration of light scattering cells on the viability and printing resolution was tested.

**Figure 4:**
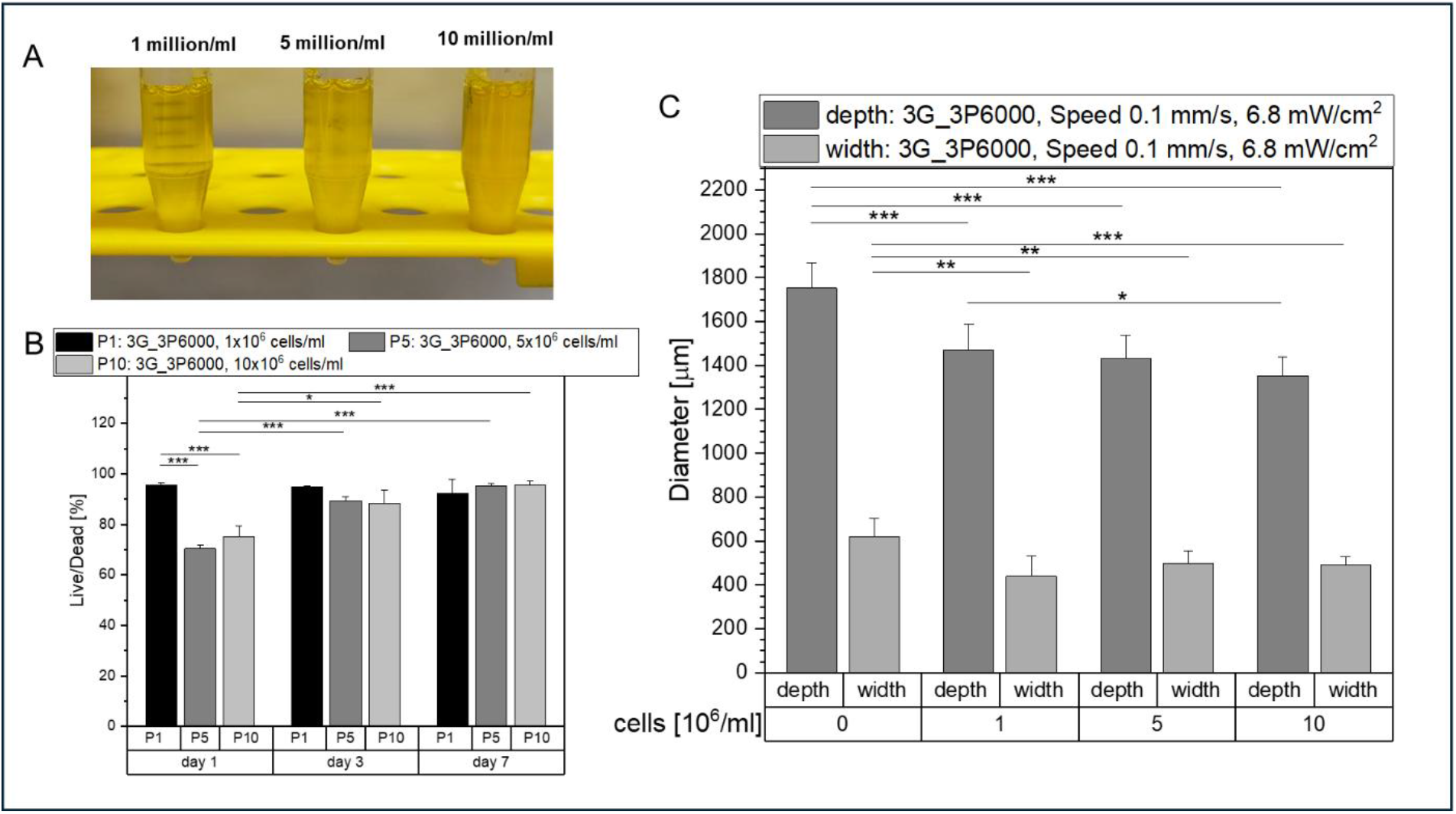
Influence of high cell concentrations (A) The 3G_3P6000 ink formulation with 1-10×10^6^ cells/ml showing an increasingly opaquer material dependent on the cell concentration demonstrated by the markers on the centrifuge tubes. (B) Live/Dead assay of the PEGDA 6000 formulations with 1×10^6^/ml, 5×10^6^/ml, and 10×10^6^/ml L929 cells for day 1-7. (C) Line width and depth of a single printed OFAB line dependent on the cells concentration, using the PEGDA 6000 formulation. The lines were measured like depicted in figure 2.

The influence of high cell concentrations on cell survival was evaluated by Live/Dead staining using the 3G_3P6000 formulation and a constant light dose for all groups. On day 1, the formulation containing 1 × 10^6^ cells mL^−1^ showed a viability of 95 ± 1%, which remained stable over 7 days. In contrast, the 5 × 10^6^ and 1 × 10^7^ cells mL^−1^ groups initially showed lower viabilities of 70 ± 1% and 75 ± 4%, respectively, but the number of living cells increased over time and reached values that were not significantly different from the 1 × 10^6^ cells mL^−1^ group by day 7. This suggests that high cell densities can be processed with OFAB under the tested conditions, although early viability may be affected by increased cellular crowding, altered diffusion, or handling-related stress. Importantly, all groups were exposed to the same nominal light dose, indicating that the observed differences were associated with formulation cellularity rather than different irradiation conditions.

The ability to process high cell densities is particularly relevant for tissue engineering applications in which native-like cellularity is required. In several musculoskeletal and cartilage-oriented biofabrication strategies, high cell concentrations are used to promote cell-cell interactions, matrix deposition, and tissue maturation. In projection-based light bioprinting, however, increasing cell density can reduce optical transparency and compromise printing accuracy due to scattering and attenuation of the projected light. In OFAB, the effect of cell-induced turbidity appears fundamentally different. Because polymerization is initiated locally at the optical fiber tip rather than through the entire resin volume, the presence of cells reduced excessive light propagation and led to a more confined curing volume under the tested conditions. This observation suggests that OFAB could be particularly useful for bioresins that are challenging for conventional light-based methods, including highly cellularized or otherwise optically turbid formulations. For a stronger mechanistic discussion, the average diameter of the L929 cells and a simple optical transmission or turbidity measurement for the different cell concentrations should be reported if available.

Overall, these data demonstrate that OFAB can process cell-laden bioresins at high cell densities while maintaining acceptable viability and improving curing confinement under the tested conditions. Rather than representing only a limitation, cell-induced turbidity may become a useful parameter for tuning the spatial resolution of optical-fiber-based bioprinting.

### 2.5 Showcasing OFABs 3D capabilities

After establishing the material and cellular processing window, we implemented a customized OFAB printer to demonstrate the fabrication of more complex 3D structures. The aim of this section is not only to highlight geometrical freedom, but also to evaluate whether single-line optimization can be translated into larger, multilayer, centimeter-scale constructs. Star-shaped structures were used as a quantifiable benchmark because their inner and outer angles, tip width, and height provide information on shape fidelity in a more realistic 3D geometry than isolated single lines.

While most of the previous research was done with a modified bioprinter using external trigger options for the light source as described by Cianciosi et. al.,^[6]^ a new custom OFAB printer was build based on a VORON filament printer. The filament dispensing head was replaced with the optical fiber and the software was modified to change the light intensity instead of the extrusion rate. This new system has an inbuild temperature control enabling constant temperatures and therefore a constant stiffness of the thermoresponsive support bath and ink at long printing times. Moreover, the printing velocity can be changed during the print and there is no more delay for turning on and off the optical fiber, which made complex prints difficult with the old system. It also enables to process STL files for OFAB taking advantage of available CAD and slicing software, which made the printing process easier compared to the system described previously. In the old system the printing of every individual line had to be programmed manually, making complex prints difficult and susceptible to errors.^[6]^ The printer in shown in Figure 5 A.

**Figure 5:**
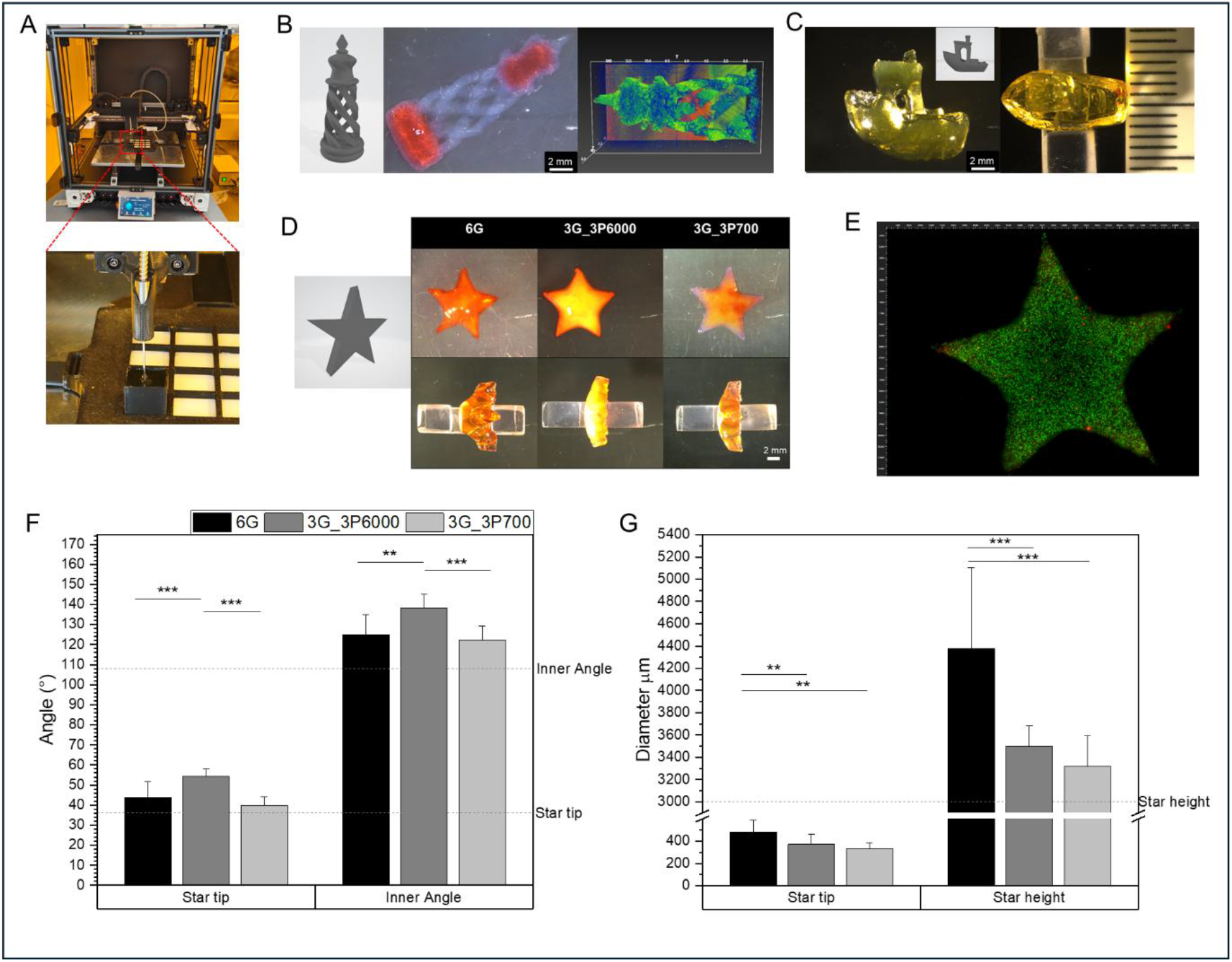
Demonstration of the custom build OFAB system based on an FFF 3d printer and demonstration of its printing abilities. (A) Picture of the custom build OFAB printer based on a VORON FFF printer. The FFF extrusion head was replaced with an optical fiber that is connected to a 405 nm LED. The build plate was modified with a Peltier element to keep the material cool during long prints. The windows were modified with an UV absorbing foil for safety. (B) The STL file, printed version, and OCT scan of a 3D OFAB printed chess figure consisted of 3% PEGDA 700 Da, 3% GelMA, 0.15% LAP and 0.015% tartrazine in 254 mM HEPES. The structure was complemented with allura red after the print to make it easier to microscope. (C) The STL file and microscope picture of an OFAB printed “benchy” consisted of 3% PEGDA 700 Da, 3% GelMA, 0.15% LAP and 0.015% tartrazine in 254 mM HEPES. (D) STL file and Microscope pictures of 3D printed stars with the different formulations at the velocity of 0.5 mm/s and an Intensity range of 6.8-5.3 mW/cm^2^. Coloured with Allura red to make the microscopy easier. (E) Microscope picture of a Live/Dead staining of a 3D printed star (3G_3P6000) consisted of 3% PEGDA 6000 Da, 3% GelMA, 0.15% LAP and 0.015% tartrazine in 254 mM HEPES with 1×10^6^ cells/ml L929 after 3 days. (F) Inner and outer angles of n = 3 stars (one of each shown in D). (G) the diameter of the tips of the stars and the hight of the stars measured on 5 positions for 3 stars.

The chess figures shown in Figure 5 B are often used as reference structures for volumetric bioprinting.^[38],[39]^ The figure was printed using the more stable 3G_3P700 formulation using the VORON based printer. This shows that OFAB can produce structures that would be commonly printed with volumetric printing. However, VBP produces resolutions down to 40 μm to 100 μm^[30],[29]^ while OFABs resolution is lower. Though the smallest line printed with OFAB, already reached a width of 70 μm at low reproducibility, OFAB produces line widths of 200-300 μm and hights of roughly 800 μm. A so-called “benchy” is often used as a benchmark for 3D printing^[20]^ and therefore was also printed with OFAB in Figure 5 C using the 3G_3P700 formulation. In Figure 5 D the different formulations were used to print a star to show how the varied materials are printable in a more complex 3D structure. Moreover, the same star was printed in Figure 5 E including 1×10^6^ /ml L929 cells and was stained after 3 days in a live dead assay, demonstrating the biocompatibility of complex 3D structures produced by OFAB. The inner angles and the angles of the tips of the 5 ponied stars from Figure 5 D were measured in Figure 5 F. The STL file had a targeted angle of 36°. While the 6G and 3G_3P700 formulations were slightly higher with 44° ± 8° and 40° ± 4° the 3G_3P6000 formulation showed the worst printing results with 54° ± 4° being significantly higher and more off target, than the other formulations. The same was observed for the inner angles which was targeted at 108°. The 6G and 3G_3P700 formulations produce angles of 125° ± 10° and 122° ± 7°. The 3G_3P6000 formulation was significantly higher with 138° ± 7°. Moreover, the overall reproducibility and homogeneity of the stars was best with 3G_3P700 and worst with G. Therefore, the better results of 6G over 3G_3P6000 might be caused by instability on in the 6G stars, since outer layers might have been rinsed away during the washing. This is possible since the printing velocity of 0.5 mm/s with I of 6.8-5.3 mW/cm^2^ reaches the limits of the 6G formulation. Moreover, the tips of the 5-pointed stars and the widths were also measured as depicted in Figure 5 G. The tips of the formulations 3G_3P700 and 3G_3P6000 with 330 ± 50 µm and 370 ± 110 µm were roughly 100 µm higher that what would be expected from a single line at 0.5 mm/s with I 6.8 mW/cm^2^ (Figure 1 G). This however can be expected when printing multiple layers on top of each other. 6G performed significantly worse than the other formulations with 480 ± 110 µm which was also seen in pervious experiments. This shows also when it comes to the width of the structures. The targeted hight of the stars was 3000 µm. The formulations 3G_3P700 and 3G_3P6000 produced hights of 3320 ± 270 µm and 3500 ± 180 µm. Since the line hight at that velocity can be assumed to be about 800 µm (Figure 1 F) and the printer was set to assume a line hight of 400 µm to assure a good overlap between the layers, those results are within expectations. The 6G formulation however was over-polymerised in the centre showing a hight of 4380 ± 730 µm with a big deviation and inhomogeneity. Overall, the 3G_3P700 formulations shows to be the best suited for high resolution printing, however the 3G_3P6000 formulations is suited to be a bio resin for 3D bioprinting. The 6G formulation with only GelMA unfortunately has relatability and resolution issues.

The 3D printing experiments demonstrate that the customized OFAB platform can translate single-line optimization into centimeter-scale hydrogel architectures. Although the current constructs still show anisotropic curing and geometry-dependent broadening, these limitations are expected for a point-wise light-based process that is still at an early stage of optimization.

The key advance is that the same platform can process both transparent and turbid formulations, can fabricate complex hydrogel structures from STL-derived toolpaths, and can maintain cell viability in printed 3D constructs. Therefore, the present results should be framed not as a final resolution benchmark, but as a demonstration that OFAB can bridge the gap between accessible embedded printing and light-based bioprinting of cell-laden hydrogels.

## 3. Conclusion

In this work, optical fiber-assisted printing was translated into a cell-compatible bioprinting approach by developing and processing GelMA- and GelMA/PEGDA-based bioresins in a freeform embedded printing configuration. The OFAB system enabled localized photocrosslinking within a hydrogel resin bath using a 405 nm LED-coupled optical fiber, while preserving the conceptual advantages of OFAP, including modularity, accessibility, and direct control over light dose through fiber velocity and irradiance.

A systematic screening of printing parameters demonstrated that both light intensity and fiber velocity strongly influence the final line geometry. Depending on the processing conditions, single-line widths in the range of 70–300 µm were obtained, while curing depth remained larger than lateral width due to the directional emission of light from the fiber tip. Among the investigated formulations, GelMA/PEGDA blends provided a favorable compromise between print resolution, construct stability, and handling after removal of the uncrosslinked bath. Rheological characterization further confirmed the relevance of GelMA thermoresponsiveness for supporting the printed structures during fabrication, while PEGDA contributed to post-crosslinking stability.

The incorporation of L929 cells established the biological feasibility of OFAB under the tested conditions. GelMA and GelMA/PEGDA 6000 Da formulations maintained high cell viability, including after printing, whereas PEGDA 700 Da-containing formulations were not suitable as bioresins despite their favorable printability. Importantly, increasing the cell concentration up to 1 × 10^7^ cells mL^−1^ did not prevent OFAB printing. Instead, the resulting turbidity reduced excessive curing and improved spatial confinement of the printed features. This finding is particularly relevant because cell-induced light scattering is commonly considered a limitation for projection-based light bioprinting methods, whereas in OFAB it may be exploited to improve curing localization.

Finally, the implementation of a customized OFAB printer enabled temperature-controlled processing and the fabrication of complex 3D hydrogel structures, including cell-laden constructs. Although further optimization is required to improve axial resolution, surface smoothness, and reproducibility across complex geometries, the present study establishes OFAB as a promising and accessible bioprinting strategy for soft, cell-laden, and optically turbid hydrogel formulations. Future developments should focus on refining fiber geometry, optimizing photoabsorber and photoinitiator concentrations, expanding the range of cytocompatible bioresins, and validating the approach with application-relevant primary cells and tissue-specific biological readouts.

## 4. Experimental selection

### Synthesis of Metacrylated-Gelatin (GelMA)

The GelMA was synthesized as described by Loessner et. al..^[40]^ 30g Gelatine (300g Bloom, porcine, Sigma Aldrich, Darmstadt, Germany) was dissolved in 300 ml distilled water. Heated to 40 °C for 2 hours and 50 °C for 10 min. 10 ml of Metacrylic anhydride (Sigma Aldrich, Darmstadt, Germany) were added and the material was shaken for 1h at 50 °C. The material was than centrifuged at 4500 rpm for 5 min. and decanted. 6000 ml destilled water were addad and the material wa transferred into 12-14k (servapor 3 dialysis tubing, MWCO 12-14k, SERVA, Heidelberg, Germany) and dialysed at 37 °C for 4 days with 8 water changes. Afterwards the material was frozen overnight at - 20°C and lyophilized for 5 days. The Degree of substitution (0.92) was determined by ^1^H-NMR using a 400 MHz Bruker Avance III HD as described by Hoch et. al.^[41]^ using the lysine content as reference as described by Zatorski et. al..^[42]^

### Synthesis of PEGDA 6000 Da

The acrylation of polyethylene glycol (PEG 2-arm, 6 kDa, Sigma Aldrich, Darmstadt, Germany) was performed for linear PEG 6000 Da as described by Türkner et. al..^[43]^ First, 50g PEG was molten at 110 °C at high vacuum for 2h. The molten PEG was dissolved in 500 ml of dry toluene (Fisher Scientific, Schwerte, Germany) in argon atmosphere. Dry triethylamine (Sigma Aldrich, Darmstadt, Germany) 3.5 ml was added, followed by addition of 3.5 ml acrylic acid. The reaction solution was stirred at room temperature for 72 h. The product was precipitated in diethyl ether (Chemobar University of Würzburg, Würzburg, Germany) two times and washed diethyl ether. The purified product was dried in vacuum at RT and obtained as white solid. The Degree of substitution (89%) was determined by ^1^H-NMR using a 400 MHz Bruker Avance III HD and a TFAA-Assay (Sigma Aldrich, Darmstadt, Germany) as previously described by Forster et. al..^[44]^

### Photoresin Preparation

The photo resin was mixed in one single step. LAP (Sigma Aldrich, Darmstadt, Germany), GelMA and if needed PEGDA 6000 Da or PEGDA 700 Da (Sigma Aldrich, Darmstadt, Germany), a 254 mM HEPES stock (Karl Roth, Karlsruhe, Germany) and a stock solution of 0.5% tartrazine (Sigma Aldrich, Darmstadt, Germany) in HEPES were added in a centrifuge tube and were shaken in a thermos shaker at 37 °C for one hour. If cells were included, they were added, and the material was shaken for another 10 min. Afterward, the material was filled in an OFAB vat and stored in the fridge for 30 min.

### OFAB 3D printing

The cooled material inside the vat was taken from the fridge and put on the individual OFAB printing platform. With the VORON based OFAB system the Peltier elements were set to 10 °C. The lines and structures were printed using a 100 nm glass optical fiber (GUL 100-BBand-SCF-custom Ferrule-SMA-3.4BX-02, N.A 0.47, Mountain Photonics, Landsberg am Lech, Germany) and a 405 nm LED (PRI Silver-LED-USB-WL, Mountain Photonics, Landsberg am Lech, Germany). After the print was finished, the vat was given in the incubator for 20 min. at 37 °C and carefully rinsed with distilled water afterwards.

### Printing resolution measurements

The single printed lines were kept in the vat, and the polymerized material was rinsed away. Afterwards, the structures were left in distilled water and scanned with Optical Computer Tomography (ATR206C1/M, Thorlabs, Bergkirchen, Germany). For the microscope pictures, the 3D prints were washed out of the vat and put in a petri dish with distilled water and then imaged with a Digital Microscope (DMS1000, Leica, Amsterdam, Netherlands).

### Rheological Measurements

The rheological measurements were conducted with Anton Paar MCR 702 rheometer (Anton Paar, Graz, Austria) with a 25 mm parallel plate geometry at a 500 µm gap. The materials were freshly prepared and measured by 5°C. The experiment included an amplitude sweep (10 rads/s) and frequency sweep (strain 1%) before and after the polymerisation and a time sweep (10 rads/s, strain 1%) during the photopolymerization. A viscosity sweep and a time sweep at (10 /1s) over the viscosity of the unpolymerized material was also conducted in a separate measurement. To polymerise the material, Photorheological measurements were performed by coupling the rheometer with a UV–vis light source (Dr. Hönle, bluepoint 4, Gräfelfing, Germany) equipped with a flexible light guide. The wavelength spectrum was narrowed using an optical filter between 390 and 500 nm. The material was polymerised for 20 seconds.

### Cell culture

Fibroblast-like cell line derived from a mouse (L929, DSMZ no.: ACC 2, murine (mouse) (Mus musculus), connective tissue fibroblasts, DMSZ, Leipzig, Germany) were cultured in Dulbecco’s Modified Eagle Medium (DMEM(1x) + GlutaMAX, gibco, Grand Island, United States) supplemented with additional 10% FCS and 1% P/S at 37 °C in 5% CO_2_ atmosphere. The cells were harvested by detaching them with 0.5% trypsin for 5 min at 37 °C, counted, centrifuged, and resuspended in the bioink. The passage 5-30 were used in the experiments. They were added to the bioresin as described previously. All prints with were conducted with a 200 µm fiber (GUL 200-BBand-SCF-custom Ferrule-SMA-3.4BX-02, N.A 0.50, Mountain Photonics, Landsberg am Lech, Germany), to make handling possible. The printed structures were detached and incubated in the Dubecco’s Modivied Eagle Medium for the required time.

### Cell viability assay

Cell viability was evaluated by counting the live and dead cells using Calcein AM and ethidium homodimer-1 (Thermo Fisher Scientific, Reinach, Switzerland) according to the manufacturer’s indications to stain the cells. Z-stack images were acquired using a fluorescence microscope (Axio Observer Colibri 7, Zeiss, Oberkochen, Germany) and processed using Fiji.

## Acknowledgements

Our research is partially financed by the European Regional Development Fund (EFRE).

## References

[1] J. Groll, T. Boland, T. Blunk, J. A. Burdick, D.-W. Cho, P. D. Dalton, B. Derby, G. Forgacs, Q. Li, V. A. Mironov, Biofabrication 2016, 8, 013001.

[2] X. Zhang, X. Zhang, Y. Li, Y. Zhang, Materials 2023, 16, 7461.

[3] J. Hagenbuchner, D. Nothdurfter, M. J. Ausserlechner, Essays in Biochemistry 2021, 65, 417–427.

[4] Y. Fang, Y. Guo, T. Liu, R. Xu, S. Mao, X. Mo, T. Zhang, L. Ouyang, Z. Xiong, W. Sun, Chinese Journal of Mechanical Engineering: Additive Manufacturing Frontiers 2022, 1, 100011.

[5] I. T. Ozbolat, W. Peng, V. Ozbolat, Drug discovery today 2016, 21, 1257–1271.

[6] A. Cianciosi, M. Pfeiffle, P. Wohlfahrt, S. Nürnberger, T. Jungst, Advanced Science 2024, 11, 2403049.

[7] C. A. Murphy, J. B. Costa, J. Silva-Correia, J. M. Oliveira, R. L. Reis, M. N. Collins, Applied materials today 2018, 12, 51–71.

[8] S. Ramesh, O. L. Harrysson, P. K. Rao, A. Tamayol, D. R. Cormier, Y. Zhang, I. V. Rivero, Bioprinting 2021, 21, e00116.

[9] D. Chimene, K. K. Lennox, R. R. Kaunas, A. K. Gaharwar, Annals of biomedical engineering 2016, 44, 2090–2102.

[10] R. Levato, K. S. Lim, W. Li, A. U. Asua, L. B. Peña, M. Wang, M. Falandt, P. N. Bernal, D. Gawlitta, Y. S. Zhang, Materials Today Bio 2021, 12, 100162.

[11] K. Zhou, Y. Sun, J. Yang, H. Mao, Z. Gu, Journal of Materials Chemistry B 2022, 10, 1897–1907.

[12] W. Hua, K. Mitchell, L. Raymond, B. Godina, D. Zhao, W. Zhou, Y. Jin, ACS biomaterials science & engineering 2021, 7, 4736–4756.

[13] D. J. Shiwarski, A. R. Hudson, J. W. Tashman, A. W. Feinberg, APL bioengineering 2021, 5.

[14] D. Nieto, J. A. Marchal Corrales, A. Jorge de Mora, L. Moroni, APL bioengineering 2020, 4.

[15] R. Levato, K. S. Lim, Biofabrication 2023, 15, 020401.

[16] J. Groll, J. A. Burdick, D.-W. Cho, B. Derby, M. Gelinsky, S. C. Heilshorn, T. Juengst, J. Malda, V. A. Mironov, K. Nakayama, Biofabrication 2019, 11, 013001.

[17] R. Rizzo, D. Ruetsche, H. Liu, M. Zenobi-Wong, Advanced Materials 2021, 33, 2102900.

[18] S. Das, B. Basu, Journal of the Indian Institute of Science 2019, 99, 405–428.

[19] R. N. Ghosh, J. Thomas, A. Janardanan, P. K. Namboothiri, M. Peter, Materials Advances 2023, 4, 5496–5529.

[20] A. P. Kitos Vasconcelos, S. C. Millik, A. Vazquez, N. Sadaba, S. Sateesh, S. Daily, S. Yu, M. Zhang, N. Manitsirisuk, A. Nelson, Annual Review of Materials Research 2025, 55, 491–521.

[21] W. Li, M. Wang, H. Ma, F. A. Chapa-Villarreal, A. O. Lobo, Y. S. Zhang, Iscience 2023, 26.

[22] S.-Y. Chang, T. Ching, M. Hashimoto, Materials Today: Proceedings 2022, 70, 179–183.

[23] B. He, J. Wang, M. Xie, M. Xu, Y. Zhang, H. Hao, X. Xing, W. Lu, Q. Han, W. Liu, Bioactive Materials 2022, 17, 234–247.

[24] B. Van Durme, A. Quaak, C. Vazquez Martel, K. Kacmaz, J. Brancart, S. Schandl, A. Ovsianikov, V. Van Rompaey, E. Blasco, Q. Thijssen, Advanced Materials Technologies 2026, e02471.

[25] X. Jing, H. Fu, B. Yu, M. Sun, L. Wang, Frontiers in Bioengineering and Biotechnology 2022, Volume 10 - 2022.

[26] A. Ovsianikov, B. N. Chichkov, in Computer-aided tissue engineering, Springer, 2012, pp. 311–325.

[27] S. Binder, F. Chalupa-Gantner, H. W. Yoo, T. Zandrini, A. Ovsianikov, Additive Manufacturing 2025, 97, 104601.

[28] Z. Faraji Rad, P. D. Prewett, G. J. Davies, Microsystems & nanoengineering 2021, 7, 71.

[29] P. N. Bernal, S. Florczak, S. Inacker, X. Kuang, J. Madrid-Wolff, M. Regehly, S. Hecht, Y. S. Zhang, C. Moser, R. Levato, Nature Reviews Materials 2025, 10, 826–841.

[30] J. Madrid-Wolff, J. Toombs, R. Rizzo, P. N. Bernal, D. Porcincula, R. Walton, B. Wang, F. Kotz-Helmer, Y. Yang, D. Kaplan, MRS communications 2023, 13, 764–785.

[31] P. N. Bernal, M. Bouwmeester, J. Madrid-Wolff, M. Falandt, S. Florczak, N. G. Rodriguez, Y. Li, G. Größbacher, R. A. Samsom, M. van Wolferen, Advanced Materials 2022, 34, 2110054.

[32] M. Pfeiffle, Master thesis, Julius-Maximilian-University Würzburg (Julius-Maximilian-University Würzburg), 2023.

[33] H. Yue, Y. Wang, S. Fernandes, C. Vyas, P. Bartolo, Macromolecular Bioscience 2025, 25, 2400587.

[34] R. Levato, O. Dudaryeva, C. E. Garciamendez-Mijares, B. E. Kirkpatrick, R. Rizzo, J. Schimelman, K. S. Anseth, S. Chen, M. Zenobi-Wong, Y. S. Zhang, Nature Reviews Methods Primers 2023, 3, 47.

[35] D. Ramos Mejia, B. Cai, S. C. Iranzo, A. Perez, Y. L. Tan, S. Lee, S. C. Heilshorn, BMC methods 2026, 3, 8.

[36] M. Fowler, A. M. Lozano, J. Krause, P. Bednarz, S. Pandey, M. Ghayour, Q. Zhang, O. Veiseh, Biomaterials Science 2025, 13, 2951–2960.

[37] S. You, Y. Xiang, H. H. Hwang, D. B. Berry, W. Kiratitanaporn, J. Guan, E. Yao, M. Tang, Z. Zhong, X. Ma, Science advances 2023, 9, eade7923.

[38] A. Cianciosi, S. Stecher, M. Löffler, P. Bauer-Kreisel, K. S. Lim, T. B. Woodfield, J. Groll, T. Blunk, T. Jungst, Advanced healthcare materials 2023, 12, 2300977.

[39] E. Krumins, J. C. Lentz, B. Sutcliffe, A. Sohaib, P. L. Jacob, B. Brugnoli, V. C. Crucitti, R. Cavanagh, R. Owen, C. Moloney, Green Chemistry 2024, 26, 1345–1355.

[40] D. Loessner, C. Meinert, E. Kaemmerer, L. C. Martine, K. Yue, P. A. Levett, T. J. Klein, F. P. Melchels, A. Khademhosseini, D. W. Hutmacher, Nature protocols 2016, 11, 727–746.

[41] E. Hoch, C. Schuh, T. Hirth, G. E. Tovar, K. Borchers, Journal of Materials Science: Materials in Medicine 2012, 23, 2607–2617.

[42] J. M. Zatorski, A. N. Montalbine, J. E. Ortiz-Cárdenas, R. R. Pompano, Analytical and bioanalytical chemistry 2020, 412, 6211–6220.

[43] E. Türker, M. S. Andrade Mier, J. Faber, S. J. Padilla Padilla, N. Murenu, P. Stahlhut, G. Lang, Z. Lamberger, J. Weigelt, N. Schaefer, Advanced Biology 2024, 8, 2400184.

[44] J. Hauptstein, L. Forster, A. Nadernezhad, H. Horder, P. Stahlhut, J. Groll, T. Blunk, J. Teßmar, Macromolecular bioscience 2022, 22, 2100331.

