## Supporting information for "Optical Fiber-Assisted Bioprinting Enables Freeform Printing of Cell-Laden and Turbid Hydrogel Resins"

**Supporting Info**

**OFAB printing steps**

The following diagram shows the single steps of OFAB.

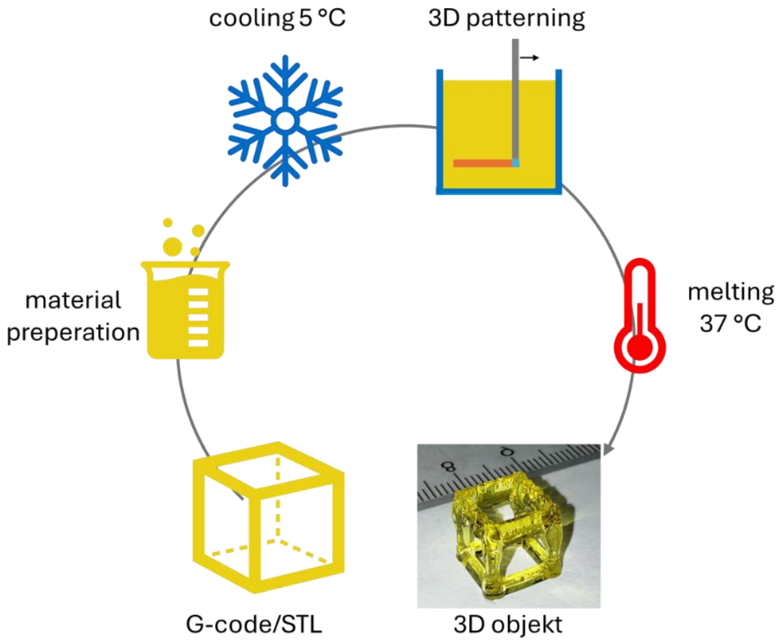


*Support Figure 1: OFAB printing steps depicted as a diagram.*

**Material formulations 3G_3P700 and 3G_3P6000**

Support Figure 2 the transparency difference of the 3G_3P700 and 3G_3P6000 formulations are shown.

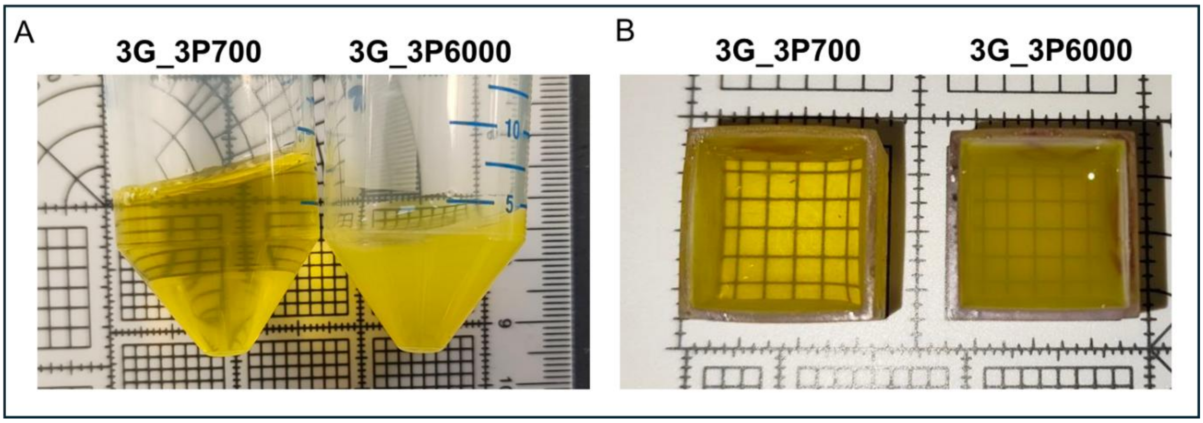


*Support Figure 2: The 3G_3P700 and 3G_3P6000 formulations using PEGDA 700 Da and PEGDA 6000 Da. The first one is fully transparent, the second one is opaque.*

### Light Intensity of the LED

Support Figure 3 shows the difference in intensity of the LED dependent on its setting.

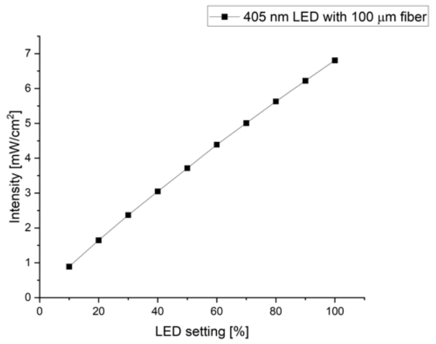


*Support Figure 3: Intensity of LED dependent on the setting.*

### Photorheologie Amplitude Sweeps

The amplitude sweeps in Support Figure 4 determined the shear strain of 1% for all following measurements, since this value was in the viscoelastic region for all materials.

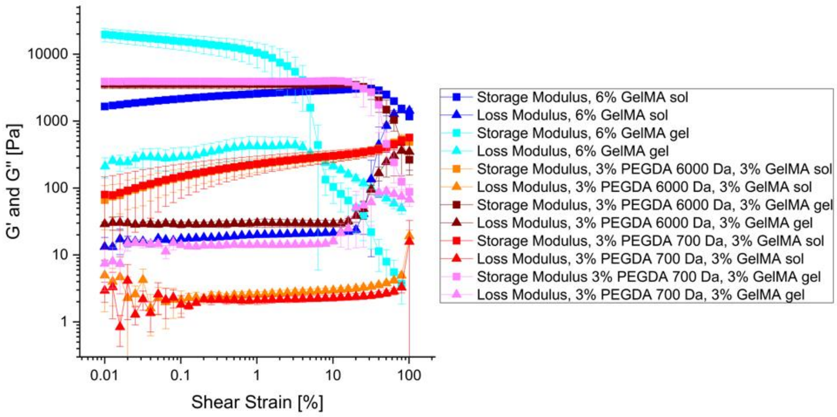


*Support Figure 4: The Amplitude sweeps before and after polymerisation for all material formulations. All include 0.015% LAP and 0.015% tartrazine in HEPES.*
